# Production of nonulosonic acids in the extracellular polymeric substances of “ *Candidatus* Accumulibacter phosphatis”

**DOI:** 10.1101/2020.11.02.365007

**Authors:** Sergio Tomás-Martínez, Hugo B.C. Kleikamp, Thomas R. Neu, Martin Pabst, David G. Weissbrodt, Mark C.M. van Loosdrecht, Yuemei Lin

## Abstract

Nonulosonic acids (NulOs) are a family of acidic carbohydrates with a nine-carbon backbone, which include different related structures, such as sialic acids. They have mainly been studied for their relevance in animal cells and pathogenic bacteria. Recently, sialic acids have been discovered as important compound in the extracellular matrix of virtually all microbial life and in “*Candidatus* Accumulibacter phosphatis”, a well-studied polyphosphate-accumulating organism, in particular. Here, bioaggregates highly enriched with these bacteria (approx. 95% based on proteomic data) were used to study the production of NulOs in an enrichment of this microorganism. Fluorescence lectin-binding analysis, enzymatic quantification, and mass spectrometry were used to analyze the different NulOs present, showing a wide distribution and variety of these carbohydrates, such as sialic acids and bacterial NulOs, in the bioaggregates. Phylogenetic analysis confirmed the potential of “*Ca*. Accumulibacter” to produce different types of NulOs. Proteomic analysis showed the ability of “*Ca*. Accumulibacter” to reutilize and reincorporate these carbohydrates. This investigation points out the importance of diverse NulOs in non-pathogenic bacteria, which are normally overlooked. Sialic acids and other NulOs should be further investigated for their role in the ecology of “*Ca*. Accumulibacter” in particular, and biofilms in general.

**Key Points:**

- “*Ca.* Accumulibacter” has the potential to produce a range of nonulosonic acids.
- Mass spectrometry and lectin binding can reveal the presence and location of nonulosonic acids.
- Role of nonulosonic acid in non-pathogenic bacteria needs to be studied in detail.

## Introduction

Wastewater transports polluting nutrients, such as organic matter, phosphorus (P) or nitrogen (N). When P and/or N are in excess, discharging this wastewater into surface waters leads to eutrophication. Thus, these pollutants must be eliminated from wastewater streams (Mainstone and Parr 2002). Enhanced biological phosphorus removal (EBPR) has become a widely applied treatment to eliminate inorganic phosphorus and organic matter from wastewater. This technology exploits the metabolic capacity of polyphosphate-accumulating organisms (PAOs) to take up inorganic phosphorus and to store it in the form of intracellular polyphosphate. “*Candidatus* Accumulibacter phosphatis”, a well-studied model PAO, has been identified as a dominant species responsible for EBPR (Seviour et al. 2003). This microorganism has not been isolated yet. It grows in the form of compact microcolonies and bioaggregates (flocs, granules or biofilms) held together by extracellular polymeric substances (EPS) (Weissbrodt et al. 2013; Barr et al. 2016).

EPS is a complex mixture of biopolymers of different nature, such as polysaccharides, proteins, nucleic acids or lipids, among others. These biopolymers are synthesized or released by microorganisms across their life cycle, forming matrices that provide mechanical stability and act as scaffold for the microorganisms in biofilms (Flemming and Wingender 2010). Although research in the past years led to analytical advances for the extraction and characterization of EPS (Felz et al. 2016; Boleij et al. 2018; Felz et al. 2019; Boleij et al. 2019), the EPS matrix still represents the “dark matter” of biofilms that need to be studied in more detail (Neu and Lawrence 2016; Neu and Lawrence 2017; Seviour et al. 2019). The pragmatic study of individual components (*i.e.* proteins and carbohydrates like monosaccharides and polysaccharides) can give new insights in the understanding of EPS as a whole. Recently, sialic acids have been detected and described in the EPS of both EBPR and salt-adapted aerobic granular sludge with the presence of “*Ca*. Accumulibacter” using fluorescence lectin-binding analysis (FLBA) coupled to confocal laser scanning microscopy (CSLM) (Weissbrodt et al. 2013; de Graaff et al. 2019).

Sialic acids are a subset of a family of α-keto acids with a nine-carbon backbone, called nonulosonic acids (NulOs). These carbohydrates are unusual among the various monosaccharide building blocks of extracellular glycoconjugates, which normally have five or six carbons. Sialic acids are typically found as terminal residues on the glycan chains of vertebrate extracellular glycoconjugates, making them the “bridging” or recognition molecules between cells, as well as between cells and extra-cellular matrix (Chen and Varki 2010). The distinct features of sialic acids contribute to higher structural complexity and the potential for more unique and varied biological functions, in comparison to other monosaccharides (Deng et al. 2013).

Looking into the specific chemical structure, sialic acids are derivatives of neuraminic (Neu) and ketodeoxynonulosonic (Kdn) acids. The most studied one is *N*-acetylneuraminic acid (Neu5Ac) (Varki et al. 2017). Apart from these acids, other NulOs have been found only in microbes, such as the isomers pseudaminic (Pse) and legionaminic (Leg) acids, which are structurally similar to sialic acids (Fig. 1A) (Knirel et al. 2003). These NulOs have recently been described as ubiquitous in the microbial world (Lewis et al. 2009; Kleikamp et al. 2020a), making their study and understanding highly relevant.

**Fig. 1.**
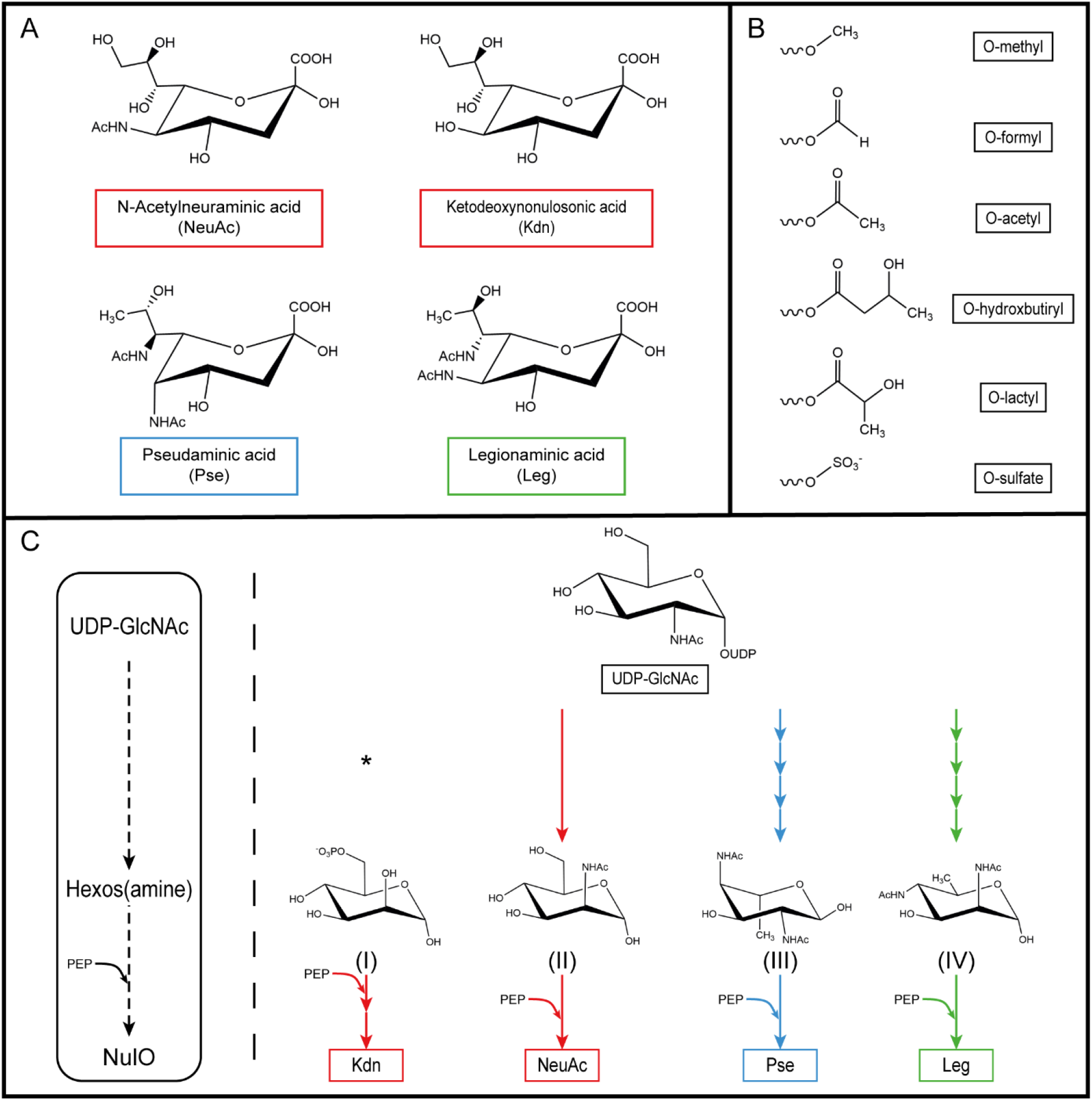
Common metabolic pathway for the biosynthesis of different NulOs. A) Chemical structure of different NulOs. B) Possible modifications of the hydroxyl groups NulOs. C) Core (left) and specific (right) biosynthetic pathways for the different NulOs. The biosynthetic pathways of the different NulOs branch from UDP-GlcNAc, with the exception of Kdn. Each arrow represents one enzymatic step. NeuAc and Kdn share the enzymes involved in the synthesis. *(I) mannose-6-phosphate; (II) N-acetylmannosamine; (III) 2,4-diacetamido-2,4,6-trideoxy-L-*altropyranose*; (IV) 2,4-diacetamido-2,4,6-trideoxy-d-mannopyranose.* Adapted from (Lewis et al. 2009)

Despite the different chemical structure, the NulOs share similarities in their metabolic pathway (Fig. 1C). The common steps in each NulO biosynthetic pathway (NAB) are catalyzed by homologous enzymes: the condensation of a 6-carbon intermediate with phosphoenol pyruvate (PEP) produces a 9-C α-keto acid (catalyzed by the enzyme NAB-2); its activation results from the addition of CMP by the enzyme NAB-1 (Lewis et al. 2009). The different NulOs can be further modified by additional substitutions on the hydroxyl groups such as *O*-acetyl, *O*-methyl, *O*-sulfate, *O*-hydroxybutiryl, *O*-formyl or *O*-lactyl groups (Fig. 1B) (Angata and Varki 2002). A phylogenetic analysis of the NAB-2 enzyme, the most conserved one in the pathway, can be used to predict the NulO types synthesized by an organism (Lewis et al. 2009). In the case of the sialic acids NeuAc and Kdn, the same biosynthetic machinery leads to their synthesis, using *N*-acetylmannosamine or mannose as starting substrate respectively (Varki et al. 2017).

Investigations of NulOs have been predominantly focused in animal cells and pathogenic bacteria. Among the large diversified NulOs, Neu5Ac is often assumed as the most dominant one present in biological samples. It plays important roles in recognition processes or stabilization of biomolecules (Hanisch et al. 2013). In animals, it is crucial to physiological processes, such as recognition between cells and neuronal transmission, or diseases such as cancer and autoimmune diseases (Traving and Schauer 1998). In pathogenic bacteria, NulOs contribute in delaying the host’s immune response by mimicking the host’s glycosylation pattern (Carlin et al. 2009).

Regardless of the intensive studies of NulOs in animal tissue and on the surface-related structure of pathogenic bacteria cells, the presence, production and function of NulOs in non-pathogenic bacteria have not been widely realized and studied. Only very recently, a genome level study (Lewis et al. 2009) and a NulOs universal survey by high resolution mass spectrometry (Kleikamp et al. 2020a) revealed the unexpectedly wide distribution of nonulosonic acid biosynthesis (NAB) pathway genes and wide-spread occurrence of NulOs in non-pathogenic bacteria. These discoveries indicate that NulOs must be an important component in the EPS of bacterial aggregates in natural and engineered ecosystems, which has been completely overlooked at present. It also indicates that the current model of evolution and utilization of sialic acids in prokaryotes which is driven by host–pathogen interactions may not reflect the complete picture and need to be questioned (Lewis et al. 2009; Kleikamp et al. 2020a).

It is known that “*Ca*. Accumulibacter” is not only the most abundant and well-studied PAO in EBPR systems, but also contributes to phosphate sequestration and phosphate cycling in estuarine systems (Watson et al. 2019). Studying the diversity, production and utilization of NulOs with an enriched culture of “*Ca*. Accumulibacter”, will add new information to the ecology of this important microorganism. Furthermore, it will provide valuable insights into the synthesis and turnover of NulOs (or sialic acids) by non-pathogenic environmental bacteria. The study of the role of NulOs outside the pathogen-host interaction will extend the current understanding of ecology and evolution of these carbohydrates.

Here, “*Ca*. Accumulibacter” was enriched using a lab-scale sequencing batch reactor with EBPR performance (Guedes da Silva 2020). NulOs produced by this biomass were analyzed by a combination of techniques, such as fluorescence lectin-binding analysis (FLBA), enzymatic release, and mass spectrometry. Genomic and proteomic investigations were conducted to evaluate the diversity of pathways involved NulOs formation and utilization by “*Ca*. Accumulibacter”.

## Materials and methods

### “*Ca.* Accumulibacter” enriched biomass and seawater-adapted aerobic granules

An in-house enrichment culture of “*Ca.* Accumulibacter” was used (Guedes da Silva 2020) was used. The enrichment was maintained in a 1.5 L sequencing batch reactor (SBR), with slight modifications from the SBR-2 described in Guedes da Silva et al. (2018). The COD-based acetate:propionate ratio in the feed was 65:35 gCOD/gCOD. FISH showed the dominance of PAO in the system (approx. 95% of biovolume), and 16S rRNA gene amplicon sequencing confirmed “*Ca.* Accumulibacter” as the dominant PAO. Proteomic investigations by (Kleikamp et al. 2020b) further confirmed this dominance (approx. 95%). Seawater-adapted aerobic granules from (de Graaff et al. 2019) were also used in the study. FISH showed the dominance of PAO in these granules as well.

### Nonulosonic acids analyses

#### Fluorescence lectin-binding analysis (FLBA)

Lectin staining of the biomass was done according to earlier works (Weissbrodt et al. 2013; Boleij et al. 2018; de Graaff et al. 2019). Bioaggregates enriched with “*Ca*. Accumulibacter” were stained and mounted in coverwell chambers with a  0.5 mm spacer in order to avoid squeezing of the samples. Glycoconjugates of the biomass were examined by means of barcoding with green fluorescent lectins (Neu and Kuhlicke 2017). Thus, all commercially available lectins labelled with a green fluorophore (FITC or Alexa488) were applied as probes individually to different aggregates. A total of 77 lectins were used to screen glycoconjugates (Bennke et al. 2013). The binding sites of the sialic acid-specific lectins that gave the strongest signal are listed in Table 1. After incubation with the lectin solution, the sample was washed with tap water for three times in order to remove unbound lectins. For 3D imaging a TCS SP5X confocal laser scanning microscope (Leica, Germany) was employed. The system comprised an upright microscope and a super continuum light source (white laser). The hardware setup was controlled by the software LAS AF 2.4.1. Confocal datasets were recorded by using 25x NA 0.95 and 63x NA 1.2 water immersion lenses. Excitation was at 490 nm and emission signals were detected simultaneously with two photomultipliers from 480 to 500 nm (reflection) and 505-580 nm (fluorescence). Image data sets were deconvolved with Huygens version 18.04 using blind deconvolution (SVI, The Netherlands) and projected with Imaris version 9.2 (Bitplane, Switzerland). Images were printed from Photoshop CS6 (Adobe).

**Table 1.**
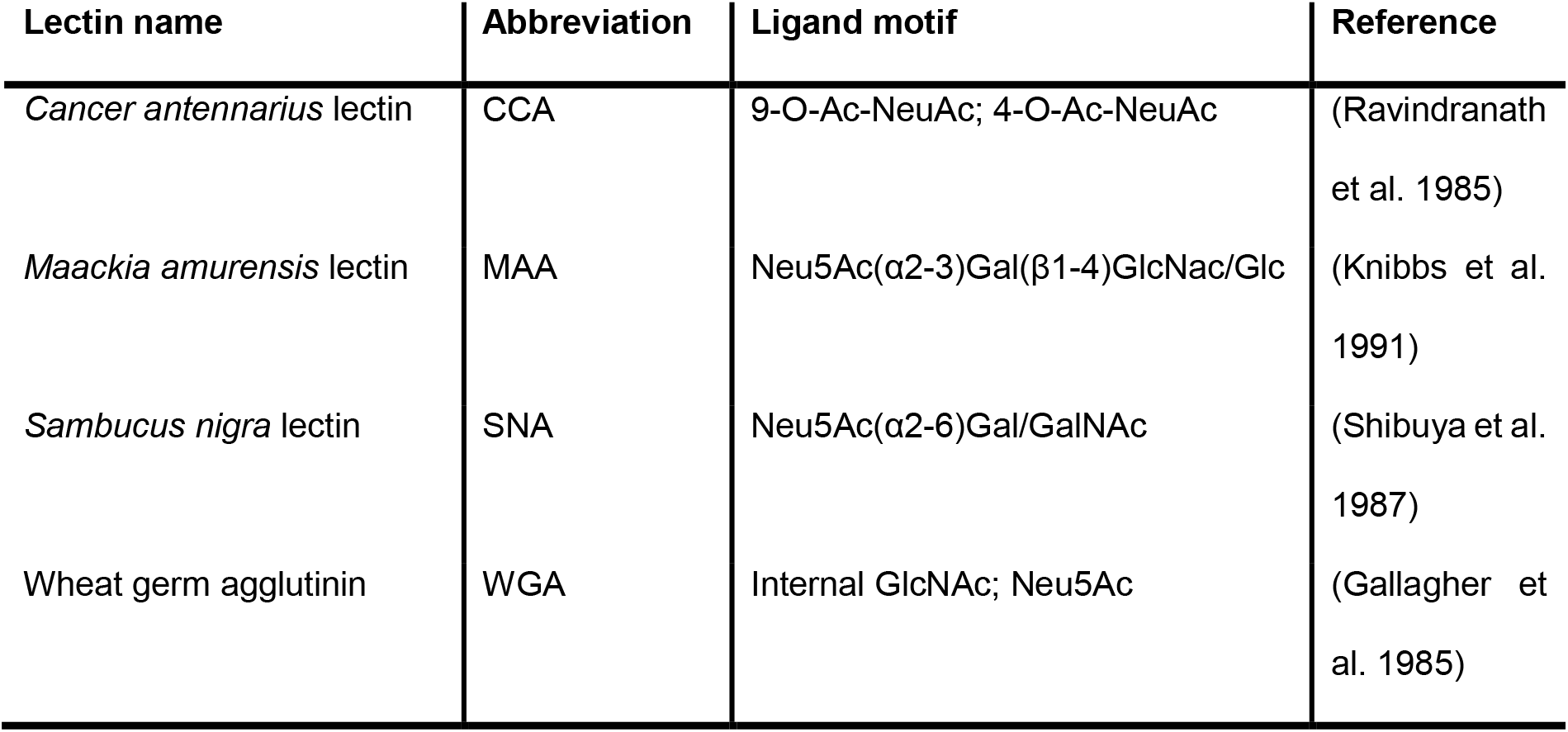
Sialic acid-specific lectin used in this analysis

| Lectin name | Abbreviation | Ligand motif | Reference |
| --- | --- | --- | --- |
| *Cancer antennarius* lectin | CCA | 9-O-Ac-NeuAc; 4-O-Ac-NeuAc | (Ravindranath et al. 1985) |
| *Maackia amurensis* lectin | MAA | Neu5Ac(α2-3)Gal(β1-4)GlcNac/Glc | (Knibbs et al. 1991) |
| *Sambucus nigra* lectin | SNA | Neu5Ac(α2-6)Gal/GalNAc | (Shibuya et al. 1987) |
| Wheat germ agglutinin | WGA | Internal GlcNAc; Neu5Ac | (Gallagher et al. 1985) |

#### Nonulosonic acid diversity and enzymatic quantification

The diversity of NulOs in bioaggregates from the “*Ca.* Accumulibacter” enrichment and in seawater-adapted aerobic granules (de Graaff et al. 2019) was analyzed during the study of Kleikamp *et al.* (Kleikamp et al. 2020a), using a high-resolution all ion reaction monitoring approach of acid hydrolyzed and DMB (1,2-diamino-4,5-methylenedioxybenzene dihydrochloride) labelled biomass. The Sialic Acid Quantitation Kit (Sigma-Aldrich, USA) was used to estimate the content of sialic acids (Neu5Ac as model one) in the enriched “*Ca*. Accumulibacter” biomass following the manual instructions. A detailed description of the protocol can be found in the Supplementary methods.

### Genomic analysis of pathways for biosynthesis of different nonulosonic acids

#### BLAST (Basic Local Alignment Search Tool) analysis of key enzymes

Different near-complete draft metagenome-assembled genomes (MAGs) of “*Ca*. Accumulibacter” (Rubio-Rincón et al. 2019) were used to get the amino acid sequences of nonulosonic acid synthases (NAB-2), *i.e.* the most conserved enzymes of the biosynthetic pathway, which condensates a 6-carbon intermediate with pyruvate to produce a 9-carbon α-keto acid (Fig. 1C). A protein sequence alignment versus a protein database (BLASTp) or a protein sequence alignment versus a translated nucleotide sequence database (TBLASTn; when only nucleotide sequences were available) from the NCBI website (blast.ncbi.nlm.nih.gov/Blast.cgi) was performed using the enzyme *N*-acetylneuraminic acid synthase from *Campylobacter jejuni* (accession number: CAL35431.1). Matches with e-value lower than 5e-15 were chosen for the phylogenetic analysis (Petit et al. 2018).

#### Phylogenetic analysis of NAB-2 sequences

In order to predict the specific NulO synthesized by the different NAB-2 enzymes of “*Ca.* Accumulibacter” the phylogenetic analysis developed by (Lewis et al. 2009) was performed. First, NAB-2 amino acid sequences (when only nucleotide sequences were available, matches were translated into amino acid sequences) from “*Ca.* Accumulibacter”, different bacteria, archaea and animals were aligned using ClustalX2 (Thompson et al. 2003), and the resulting nexus files were uploaded into PAUP* 4.0a166 (Swofford 2001). Gaps and domains not found in all sequences were excluded and a neighbor-joining tree was constructed using the bootstrap/jackknife option with 1,000 replicates. Less conserved enzymes from “*Ca.* Accumulibacter” were removed from the analysis in order to improve the final alignment. The removed sequences might represent unknown specificities not included in the analysis (Lewis et al. 2009).

### Shotgun proteomics analysis

Shotgun proteomic analysis of the “*Ca.* Accumulibacter” enrichment was performed as described in the study of (Kleikamp et al. 2020b). A detailed description of the methodology can be found in the Supplementary methods.

## Results

The enrichment culture of “*Ca.* Accumulibacter” was derived from the system described by Guedes da Silva (2020). Data describing the performance of the enrichment are given by the authors. As shown by fluorescence *in situ* hybridization (FISH) and proteomic data (Kleikamp et al. 2020b), the bioaggregates used in this research were highly enriched with “*Ca*. Accumulibacter” (approx. 95%).

### Nonulosonic acid analyses

#### Fluorescence lectin-binding analysis (FLBA)

Lectins are proteins that bind to specific carbohydrate groups. Fluorescence-labelled lectins can be used as probes for the *in situ* analysis of glycoconjugates in the EPS of bioaggregates (Neu and Kuhlicke 2017). Intact biomass samples collected from the “*Ca*. Accumulibacter” enrichment culture were screened with 77 lectins (data not shown). Some sialic acid-specific lectins gave a positive result, such as CCA, WGA, MAA and SNA (the binding site of these lectins can be found in Table 1). Especially, MAA and SNA gave the strongest signal (Fig. 2A and B). The signals from both lectins were widely distributed across the aggregate, mainly detected at the surface of the bacterial cells (A more detailed view can be seen in Fig. S 1). SNA recognizes α-2,6-linked sialic acid; while MAA recognize α-2,3-linked sialic acid (Soares et al. 2000) (Table 1). The strong signals from both SNA and MAA lectins indicated that the sialic acids on the cell surface of “*Ca*. Accumulibacter” present both types of linkages. In contrast with SNA and MAA, the distribution of the signal of WGA showed the presence of lectin-specific glycoconjugates in other parts of the biomass (Fig. 2C). This may be due to the wider specificity of WGA (*e.g.* towards GlcNAc). In addition, although CCA gave a low signal (Fig. 2D), it indicated the presence of sialic acids with different modifications, as it is specific for the staining of 9-O or 4-O-acetyl NeuAc.

**Fig. 2.**
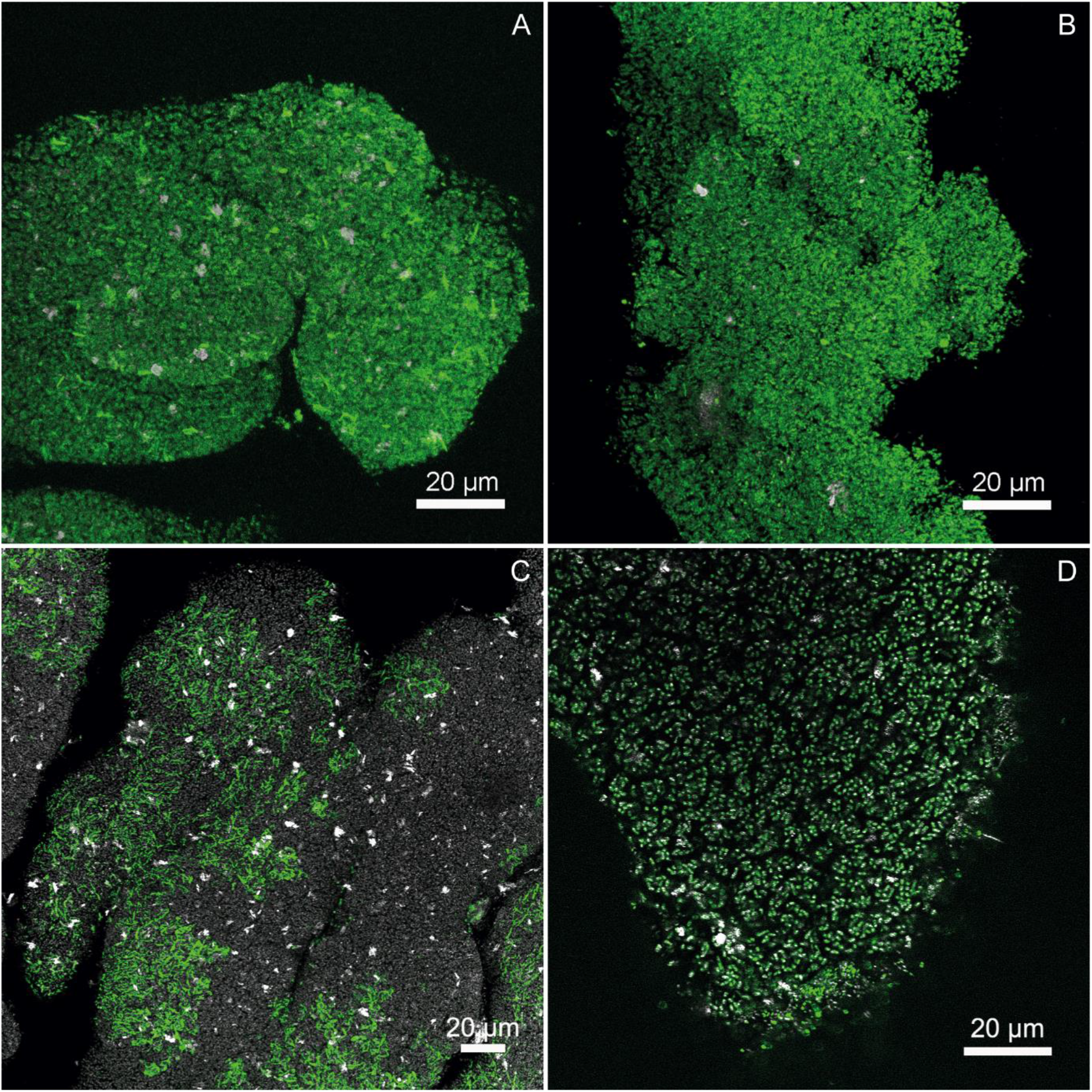
Confocal laser scanning microscopy (CLSM) after fluorescence lectin-binding analysis (FLBA). Images show bioaggregates enriched in “*Ca*. Accumulibacter”. The glycoconjugates visualized in A, B, C, D show four different sialic acid-specific lectins (A - MAA, B - SNA, C - WGA, D - CCA). The reflection signals either mark reflective particles associated with the granules (A, B) or outline the shape of the granule due to cell internal reflections (C, D). Color allocation: green – glycoconjugates, white – reflection signal

#### Nonulosonic acid diversity by mass spectrometry analysis

Lectin staining showed a wide distribution of different sialic acids in the “*Ca*. Accumulibacter” enriched bioaggregates. However for this, not only the type of NulO but also the same motifs (linkage type and subterminal monosaccharide) can result in binding. A recent molecular survey by Kleikamp et al. (2020a) demonstrated the diversity of NulOs across many species, including “*Ca*. Accumulibacter” and seawater-adapted aerobic granules dominated with “*Ca*. Accumulibacter” (de Graaff et al. 2019). Where bacterial NulOs (Pse/Leg) were dominant in both samples, also sialic acids were present in smaller quantities. Small amounts of Kdn, in the case of “*Ca*. Accumulibacter”, and NeuAc, for seawater-adapted aerobic granules, could be detected.

#### Sialic acid enzymatic release

Bacterial NulOs and sialic acids are widely distributed in the enriched “*Ca*. Accumulibacter” biomass. In order to quantify the amount of sialic acids present in the enrichment, a commercial enzymatic assay was performed, using a broad spectrum sialidase (α-(2→3,6,8,9)-neuraminidase). The enzyme releases α-2,3-, α-2,6-, α-2,8-, and α-2,9-linked NeuAc. The liberated sialic acids are then detected and quantified after a reaction with an aldolase and dehydrogenase. Unfortunately, no sialic acids were released. This could be due to the specificity of the sialidase which might recognize only NeuAc, but not other variants of sialic acids (*e.g.* Kdn) and bacterial NulOs. According to MS analysis, NeuAc was present in seawater-adapted aerobic granules, but not in the enriched “*Ca.* Accumulibacter” biomass, which explains why the amount of sialic acids was successfully quantified by the enzymatic assay in seawater-adapted granules as described in de Graaff et al. (2019), but was unsuccessful for “*Ca.* Accumulibacter” enrichment. In fact, sialidases have been reported to differ in their sensitivity, *e.g.* it was found that Kdn is linked to almost all glycan structures in place of NeuAc, but it has lower sensitivity to sialidase which is specific for NeuAc (Lambre et al. 1982).

### Phylogenetic analysis

Various metagenome-assembled genomes (MAGs) of “*Ca*. Accumulibacter” available in public repositories and surveyed in literature (Rubio-Rincón et al. 2019) were used to predict their potential to produce NulOs and their possible diversity. The prediction was focused on the NulO synthase (NAB-2), the enzyme that condenses a 6-carbon intermediate with phosphoenolpyruvate to yield a 9-carbon α-keto acid. This is a common step in the biosynthetic pathway of the different NulOs and the most conserved enzyme in the metabolic route (Lewis et al. 2009). The enzyme NeuAc synthase from *Campylobacter jejuni* (accession number: CAL35431.1) was used to obtain NAB-2 amino acid sequences from the different genomes of “*Ca.* Accumulibacter” (Table 2). Different reported NAB-2 amino acid sequences were used as query, giving similar results (data not shown). All the potential NAB-2 enzymes listed present low e-value, ranging from 6e-53 to 5e-19. Some available genomes did not show the presence of this enzyme, which can be both due to a poor assembly or annotation of the genome or to the genetic incapacity of some genotypes to produce NulOs.

**Table 2.**
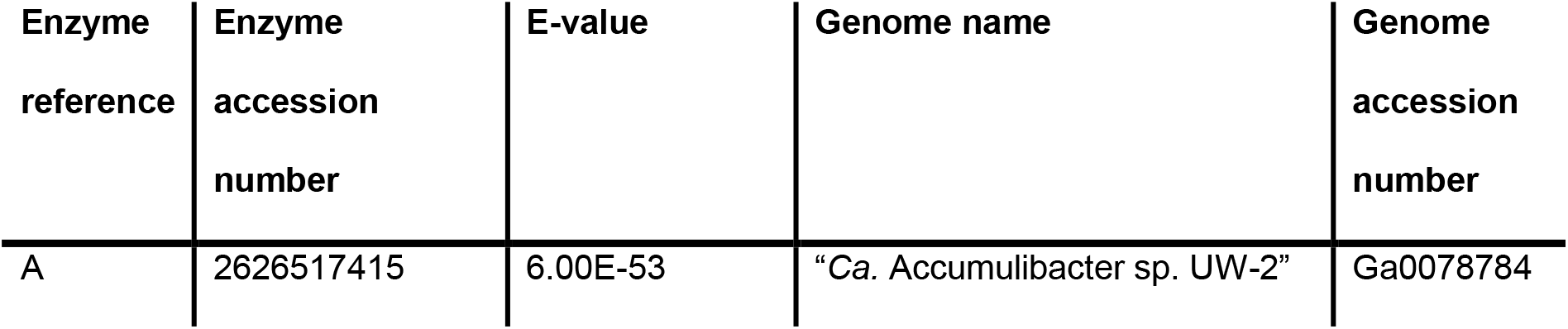
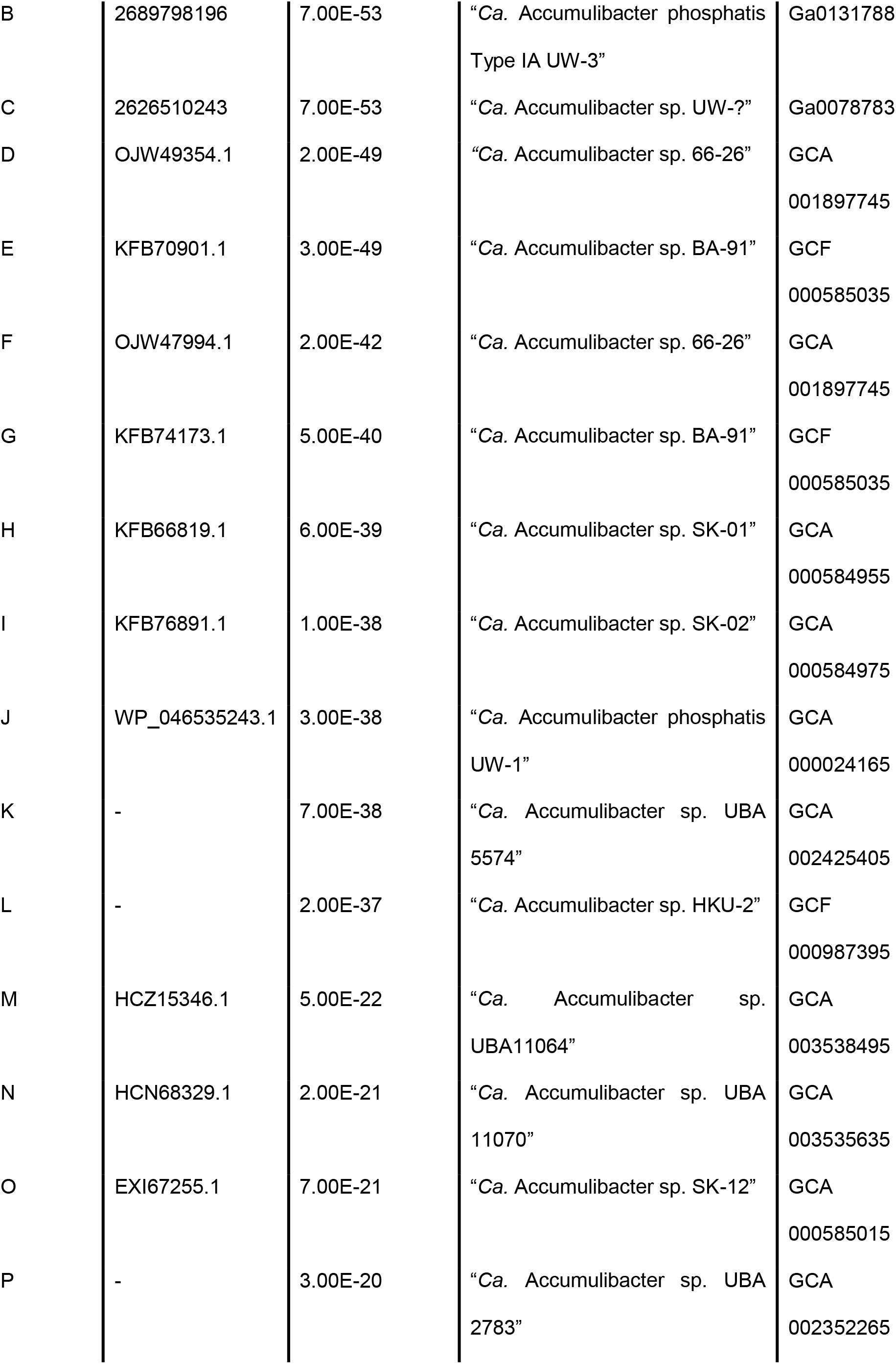
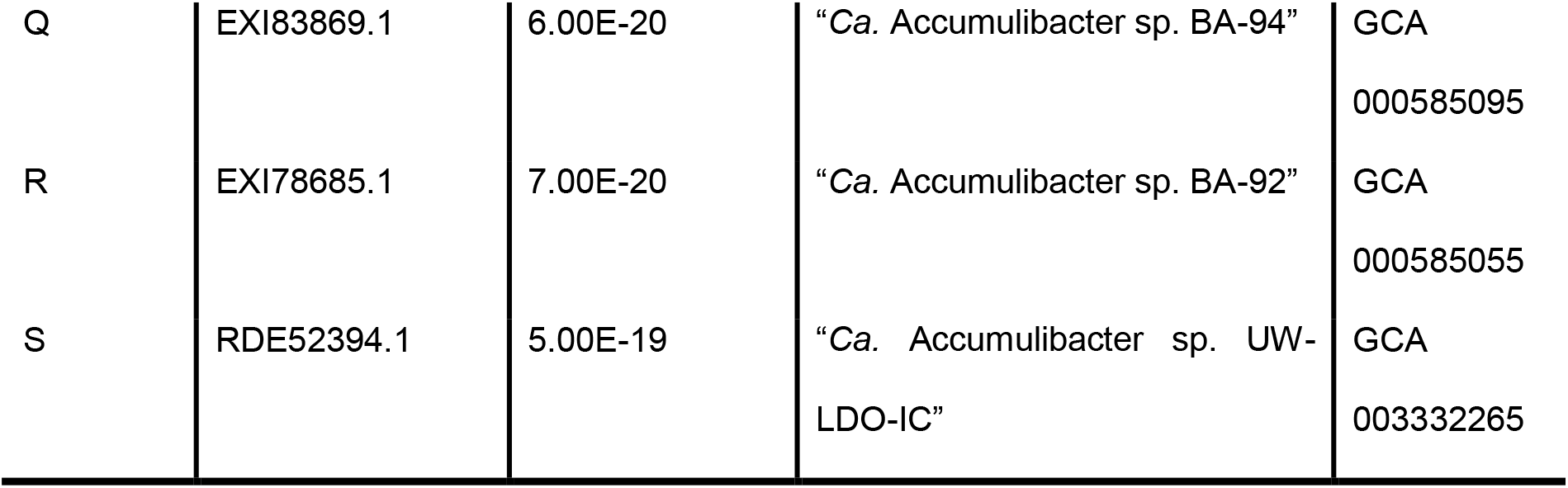
Selected NAB-2 enzymes (nonulosonic acid synthase) from “*Ca.* Accumulibacter” used for the phylogenetic analysis. Amino acid sequences were obtained by performing BLASTp using the NAB-2 enzyme from *C. jejuni* (accession number: CAL35431.1) as query. No accession numbers are provided for the enzymes where no protein sequences were available as the whole nucleotide sequence was used. Genomes were recovered from on-line public databases such as NIH GenBank (*GCA accession numbers*), NCBI RefSeq (*GCF accession numbers*), and JCI MGM (*Ga accession numbers*)

These amino acid sequences were used to predict their potential specificity using the phylogenetic analysis method developed by Lewis *et al*., (Lewis et al. 2009). The sequences of “*Ca*. Accumulibacter” with the higher e-values (m-s in Table 2) were eliminated from the analysis as they appeared to be less-conserved and affected the multiple alignment and therefore, the phylogenetic analysis. These divergent sequences might indicate specificity for a different NulO than the ones used for the final analysis (Lewis et al. 2009). The rest of enzymes of “*Ca*. Accumulibacter”, together with sequences from animal, bacteria and archaea, were used to generate a distance-based neighbor-joining tree (Fig. 3).

**Fig. 3.**
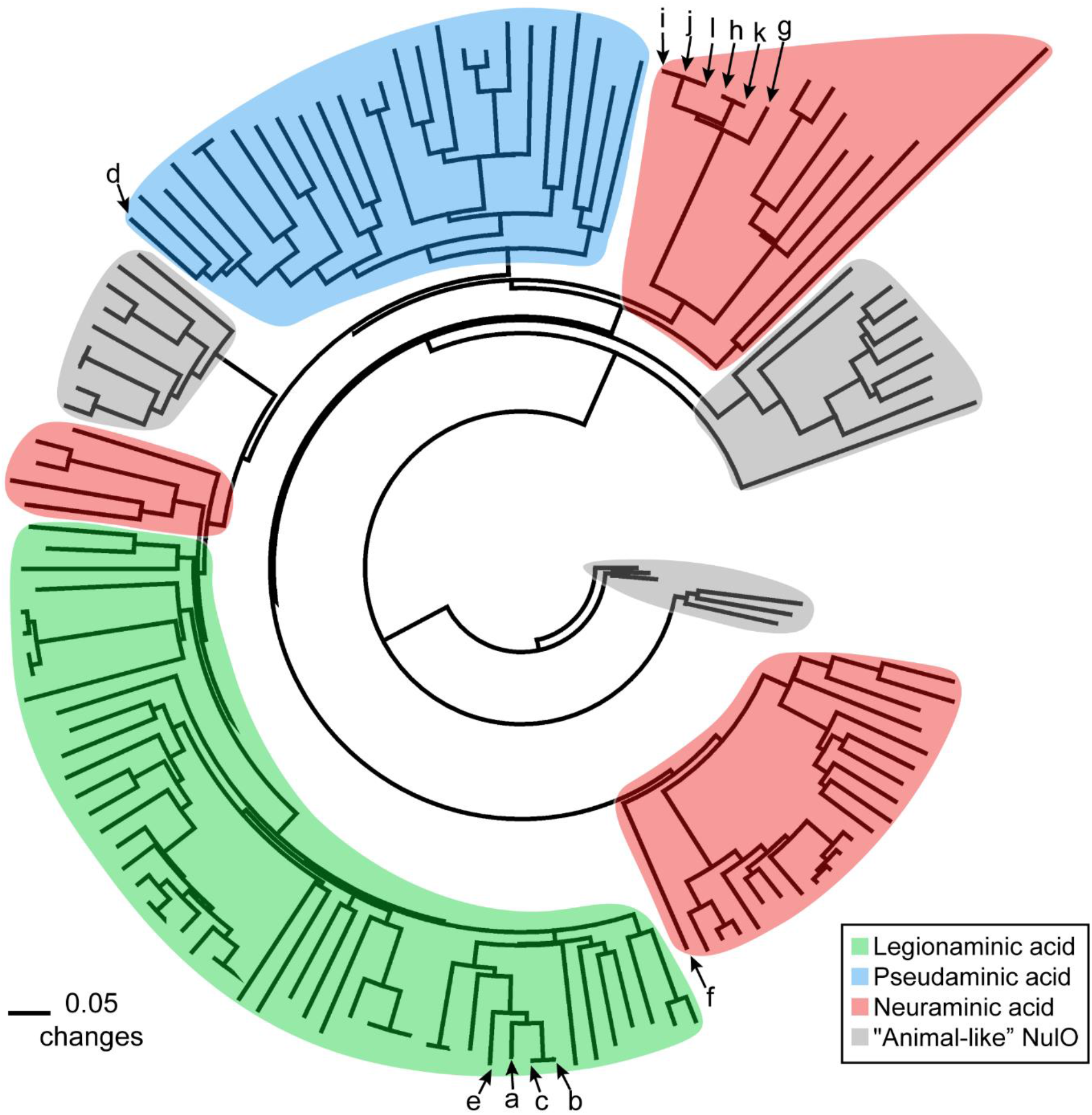
Distance-based neighbor-joining tree of the NAB-2 sequences (nonulosonic acid synthase). Sequences from bacteria, archaea and animals were used. Enzymes are grouped based on their predicted nonulosonic acid specificity (color shading). Letters (a-l) indicate the enzymes present in the different available genomes of “*Ca*. Accumulibacter” as shown in Table 2

The different NAB-2 sequences were grouped based on their predicted specificities. Four groups are generated corresponding to legionaminic acid (Leg), pseudaminic acid (Pse), neuraminic acid (Neu) and “animal-like” NulOs, which reflects a novel phylogenetic class for which no biochemical data currently exist (Lewis et al. 2009). The NAB-2 enzymes obtained from the genomes of “*Ca*. Accumulibacter” reported in literature were present in different NulOs groups with proven biochemical data (*i.e.* Pse, Leg and Neu). In two genomes, “*Ca*. Accumulibacter sp. BA-91” and “*Ca*. Accumulibacter sp. 66-26”, two different NAB-2 sequences were found in each genome. In both cases, the predicted synthesized NulO by each enzyme copy was different, indicating “*Ca*. Accumulibacter sp. BA-91” can synthesize both Leg and Neu, and “*Ca*. Accumulibacter sp. 66-26” can produce both Pse acid and Neu. Therefore, it is predicted that “*Ca*. Accumulibacter” has the potential to synthesize NulOs with one or two different core structures (Pse/Leg and Neu). Looking back at the mass spectrometry (MS) results, the enriched “*Ca*. Accumulibacter” in this research produced bacterial NulOs (as Leg and Pse are isomers, they cannot be differentiated by MS analysis) and Neu (including Kdn and derived forms of Neu (*e.g.* NeuAc) as well since they share the same pathway with Neu), which is in consistence with the phylogenetic prediction.

### Interpretation of proteomic analysis

The production of NulOs by “*Ca*. Accumulibacter” was confirmed by lectin staining, the recent mass spectrometry survey (Kleikamp et al. 2020a) and the phylogenetic analysis. To understand also the metabolism of NulOs, the proteome of the “*Ca*. Accumulibacter” enriched biomass was studied using mass spectrometry. Out of the complete list of identified proteins (Table S 1), it was found that “*Ca.* Accumulibacter” expressed neuraminic acid receptor, permease and tripartite ATP-independent periplasmic (TRAP) transporter proteins (Table 3). Those three proteins are used by some pathogenic bacteria as mechanism to decorate their surface molecules, such as capsule polysaccharides, lipopolysaccharides or flagellum, with sialic acids scavenged from the host. An extracellular sialic acid molecule is captured via a receptor (*i.e.* neuraminic acid receptor) and transported through the plasma membrane into the cell via a transporter (*e.g.* TRAP transporter); linking the sialic acid to a glycoconjugate and finally embedding the glycoconjugate within the plasma membrane (Honma et al. 2015). This suggests the presence of a NulOs-specific utilization/recycling system in the “*Ca*. Accumulibacter” enrichment, similar to the one in pathogenic bacteria. Other enzymes involved in the utilization of NulOs were not found, such as sialidases or enzymes involved in the catabolism of NulOs.

**Table 3.**
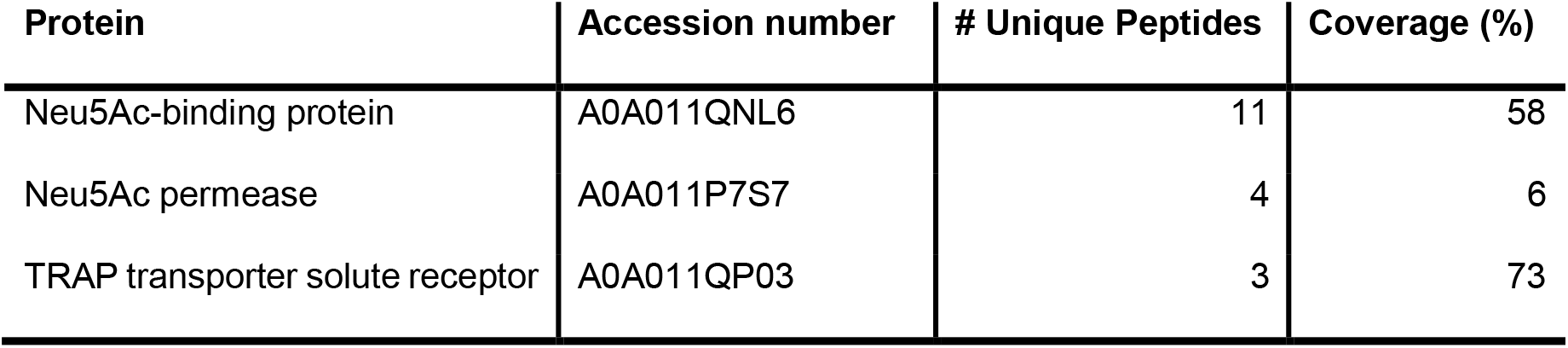
Expressed proteins by “*Ca*. Accumulibacter” involved in the transport of NulOs. The complete list of identified proteins in the enriched biomass can be found in Table S 1

| Protein | Accession number | # Unique Peptides | Coverage (%) |
| --- | --- | --- | --- |
| Neu5Ac-binding protein | A0A011QNL6 | 11 | 58 |
| Neu5Ac permease | A0A011P7S7 | 4 | 6 |
| TRAP transporter solute receptor | A0A011QP03 | 3 | 73 |

## Discussion

### “*Ca*. Accumulibacter” and nonulosonic acid production

Nonulosonic acids (NulOs) are a family of acidic carbohydrates with a nine-carbon backbone. They include sialic acids and other bacterial monosaccharides such as pseudaminic (Pse) and legionaminic acids (Leg). NulOs have been observed at the surface of animal cells and pathogenic bacteria, but they have been generally overlooked in non-pathogenic microorganisms (Varki et al. 2017).

Recently, sialic acids were discovered in glycoproteins within the extracellular polymeric substances (EPS) of seawater-adapted aerobic granular sludge (de Graaff et al. 2019). “*Ca*. Accumulibacter” was suggested to be responsible for sialic acid production. However, to prove the link between the specific microorganism and sialic acid production, a study on a highly enriched culture was necessary. The granular biomass used in this study was proven to be highly enriched in “*Ca*. Accumulibacter” (approx. 95%, by proteomic investigations (Kleikamp et al. 2020b) and by FISH staining (Guedes da Silva et al. 2018)). Sialic acid-specific lectin staining displayed that sialic acids with α-2,3- and α-2,6-linkage to the sub-terminal monosaccharide were distributed widely on the cell surface of “*Ca*. Accumulibacter”. In fact, these sialic acids visualized by lectin staining consist of diverse NulOs: *i.e.* Kdn (a common sialic acid) and Pse/Leg (bacterial NulOs) with various modifications, as the actual structure cannot be determined by lectin staining (Song et al. 2011). The most conserved enzyme (nonulosonic acid synthase, NAB-2) of the NulOs biosynthetic pathway can be traced back from the available genomes of “*Ca*. Accumulibacter”. The lack of this enzyme in some genomes might be attributed to a low quality or an incomplete state of those genomes but also to the genetic inability to produce NulOs of some genotypes of “*Ca*. Accumulibacter”. Phylogenetic analysis based on different sequences of the NAB-2 enzyme predicted the capacity of “*Ca*. Accumulibacter” to produce Pse, Leg and/or Neu (including Kdn). Therefore, this study provides evidence that “*Ca.* Accumulibacter” can synthesize sialic acids and other NulOs. Moreover, it shows the significant diversity of NulOs available in biological environments, in addition to the most common sialic acid Neu5Ac.

### Importance of sialic acids and other nonulosonic acids in non-pathogenic bacteria

The ability of bacteria to synthesize sialic acids has been mainly studied in a number of pathogens, where sialic acids or NulOs serve as a way of abolishing the immune response of the host by molecular mimicry [19]. Three different types of NulOs are frequently reported as produced by pathogenic bacteria: NeuAc, Pse and Leg. Most NulOs producing pathogens synthesize one type of NulOs, *e.g. Pasteurella multocida* can synthesize NeuAc, *Pseudomonas aeruginosa* can synthesize Pse, and *Clostridium botulinum* can synthesize Leg. Only few pathogens, such as *Campylobacter jejuni* and *Vibrio vulnificus* can synthesize multiple types of NulOs depending on the strain examined (Almagro-Moreno and Boyd 2010). The production of NulOs confers specific advantages to these bacteria in the host-pathogen interaction. Surprisingly, “*Ca.* Accumulibacter”, a non-pathogenic bacterium that was cultivated in a bioreactor without any host-pathogen interaction, were found to produce different types of NulOs. Moreover, these compounds were present in most bacteria and archaea recently tested by Kleikamp *et al.* (Kleikamp et al. 2020a). Thus, the synthesis of NulOs is not necessarily connected with the host-pathogen interaction.

Apart from the synthesis of NulOs, mammalian commensal and pathogenic bacteria that colonize sialic acid rich tissues, such as the respiratory or the gastrointestinal tract, use host-derived sialic acids as competitive advantage. These bacteria take up sialic acids released from the host by means of dedicated transporters, either incorporating them into their cell surface macromolecules or metabolizing them as a source of carbon, nitrogen and energy source (Almagro-Moreno and Boyd 2010). These bacteria that uptake/utilize sialic acids are closely associated with the host and exposed to a sialic acids rich environment. However, it is extremely interesting to see that “*Ca.* Accumulibacter”, cultivated without any NulOs in the media, still expressed Neu receptor, permease and TRAP transporter proteins, which are ssential for the uptake of NulOs. Therefore, the common understanding that both the abilities to synthesize and utilize NulOs are limited within pathogenic and/or commensal bacteria is not correct. These abilities might be widely spread in bacteria.

Most of the studies of NulOs to date have been focused on Neu5Ac, since it is the most abundant one in mammals (especially in humans). The findings that there were multiple NulOs produced by “*Ca.* Accumulibacter” enrichment and microorganisms in seawater-adapted aerobic granules, together with other findings reported in literature, suggest the need to extend the consideration of NulOs beyond Neu5Ac alone when bacteria are involved.

### Nonulosonic acids and the EBPR process

To avoid eutrophication due to phosphorus pollution, inorganic phosphorus is removed from wastewater by a process called EBPR. “*Ca.* Accumulibacter” has been identified as the dominant organisms responsible for EBPR (Zilles et al. 2002). This microorganism has been well studied in the past decades: different genomic, proteomic, metabolic and modelling studies are available (Oehmen et al. 2010; Barr et al. 2016; Oyserman et al. 2016; Guedes da Silva et al. 2018; Guedes da Silva et al. 2019; Rubio-Rincón et al. 2019). However, most of the studies overlook the extracellular matrix.

Sialic acids are known to participate in pathogen-cell interaction and intracellular communication processes in mammals (Schnaar et al. 2014). NulOs might play a similar role in “*Ca*. Accumulibacter” and provide advantages in competing with other microorganisms in the EBPR process. “*Ca*. Accumulibacter” is able to synthesize multiple NulOs with various modifications. These structures cannot be recognized by a single type of sialidase (as shown in the enzymatic analysis), therefore they can protect the cells from enzymatic degradation. On the other hand, when NulOs become available (*e.g.* released from their own macromolecules), the expression of specific transporters by “*Ca.* Accumulibacter” allows them to re-uptake these carbohydrates and re-utilize them, avoiding synthesizing them *de novo*. Through this recycling, less nutrients and cellular energy resources are required. Moreover, in vertebrates, sialic acids are typically found as terminal residues on the glycan chains of extracellular glycoconjugates, acting as “bridging” molecules between cells, and between cells and extracellular matrices (Chen and Varki 2010). At this point, NulOs may be involved in regulating “*Ca.* Accumulibacter” bioaggregates formation in wastewater treatment process and natural estuarine systems; they could also act as recognition sites for bacteriophages. Therefore, sialic acids and other nonulosonic acids should be investigated in further detail to understand their role in the ecology of “*Ca.* Accumulibacter” and even in the EBPR process in particular, and biofilms in general.

## Declarations

### Ethical approval

Not applicable

### Consent to participate

Not applicable

### Consent for publication

Not applicable

### Availability of data and material

The data generated and/or analyzed during the current study are included in this article and its supplementary material.

### Code availability

Not applicable

### Authors’ contributions

STM and YL planned the research based on intensive discussions among all the authors, especially with MvL and DW. MP and HK conducted the mass spectrometry analyses. TN performed the fluorescence lectin-binding analysis. STM worked on the phylogenetic analysis and additional lab work. STM and YL interpreted the data and played major roles in drafting, writing and revising the manuscript. All authors read and approved the manuscript.

## Acknowledgements

The authors would like to thank Leonor Guedes da Silva and Danny de Graaff for providing the biomass used in these investigations, and Ute Kuhlicke for her help with the FLBA-CSLM analyses and digital images processing.

## Supplementary information

### Supplementary methods

Shotgun proteomic analysis and Enzymatic quantification.

**Table S 1** Complete list of unique proteins identified in the enrichment of “*Ca.* Accumulibacter”

**Fig. S 1** Zoomed in section of a granule showing more details of glycoconjugate distribution after lectin staining. The settings for projections were defined in a way that strong and weak signal can be differentiated in different colors. A) Strong signal of lectin MAA at the surface of the bacterial cells (shown in green). Please take notice of the blob-like appearance of the glycoconjugates at the bacterial cell surface. B) Weak signal of the lectin MAA in the space in between bacterial cells (shown in red). C) Overlay of A and B. In Figure D), E) and F) the same is shown for the lectin SNA

## References

Almagro-Moreno S, Boyd EF (2010) Bacterial catabolism of nonulosonic (sialic) acid and fitness in the gut. Gut Microbes 1:45–50. https://doi.org/10.4161/gmic.1.1.10386

Angata T, Varki A (2002) Chemical diversity in the sialic acids and related α-keto acids: An evolutionary perspective. Chem Rev 102:439–469. https://doi.org/10.1021/cr000407m

Barr JJ, Dutilh BE, Skennerton CT, Fukushima T, Hastie ML, Gorman JJ, Tyson GW, Bond PL (2016) Metagenomic and metaproteomic analyses of Accumulibacter phosphatis-enriched floccular and granular biofilm. Environ Microbiol 18:273–287. https://doi.org/10.1111/1462-2920.13019

Bennke CM, Neu TR, Fuchs BM, Amann R (2013) Mapping glycoconjugate-mediated interactions of marine Bacteroidetes with diatoms. Syst Appl Microbiol 36:417–425. https://doi.org/10.1016/j.syapm.2013.05.002

Boleij M, Pabst M, Neu TR, van Loosdrecht MCM, Lin Y (2018) Identification of Glycoproteins Isolated from Extracellular Polymeric Substances of Full-Scale Anammox Granular Sludge. Environ Sci Technol 52:13127–13135. https://doi.org/10.1021/acs.est.8b03180

Boleij M, Seviour T, Wong LL, van Loosdrecht MCM, Lin Y (2019) Solubilization and characterization of extracellular proteins from anammox granular sludge. Water Res 164:114952. https://doi.org/10.1016/J.WATRES.2019.114952

Carlin AF, Uchiyama S, Chang YC, Lewis AL, Nizet V, Varki A (2009) Molecular mimicry of host sialylated glycans allows a bacterial pathogen to engage neutrophil Siglec-9 and dampen the innate immune response. Blood 113:3333–3336. https://doi.org/10.1182/blood-2008-11-187302

Chen X, Varki A (2010) Advances in the biology and chemistry of sialic acids. ACS Chem. Biol. 5:163–176

de Graaff DR, Felz S, Neu TR, Pronk M, van Loosdrecht MCM, Lin Y (2019) Sialic acids in the extracellular polymeric substances of seawater-adapted aerobic granular sludge. Water Res 155:343–351. https://doi.org/10.1016/J.WATRES.2019.02.040

Deng L, Chen X, Varki A (2013) Exploration of sialic acid diversity and biology using sialoglycan microarrays. Biopolymers 99:650–665

Felz S, Al-Zuhairy S, Aarstad OA, van Loosdrecht MCM, Lin YM (2016) Extraction of Structural Extracellular Polymeric Substances from Aerobic Granular Sludge. J Vis Exp e54534. https://doi.org/10.3791/54534

Felz S, Vermeulen P, van Loosdrecht MCM, Lin YM (2019) Chemical characterization methods for the analysis of structural extracellular polymeric substances (EPS). Water Res 157:201–208. https://doi.org/10.1016/J.WATRES.2019.03.068

Flemming H-C, Wingender J (2010) The biofilm matrix. Nat Rev Microbiol 8:623–633. https://doi.org/10.1038/nrmicro2415

Gallagher JT, Morris A, Dexter TM (1985) Identification of two binding sites for wheat-germ agglutinin on polylactosamine-type oligosaccharides. Biochem J 231:115–122. https://doi.org/10.1042/bj2310115

Guedes da Silva L (2020) Life in changing environments: The intriguing cycles of Polyphosphate Accumulating Organisms. Dissertation. Delft University of Technology

Guedes da Silva L, Gamez KO, Gomes JC, Akkermans K, Welles L, Abbas B, van Loosdrecht MCM, Wahl SA (2018) Revealing metabolic flexibility of Candidatus Accumulibacter phosphatis through redox cofactor analysis and metabolic network modeling. bioRxiv 458331. https://doi.org/10.1101/458331

Guedes da Silva L, Tomás-Martínez S, van Loosdrecht MCM, Wahl SA (2019) The environment selects: Modeling energy allocation in microbial communities under dynamic environments. bioRxiv 689174. https://doi.org/10.1101/689174

Hanisch F, Weidemann W, Großmann M, Joshi PR, Holzhausen HJ, Stoltenburg G, Weis J, Zierz S, Horstkorte R (2013) Sialylation and muscle performance: Sialic acid is a marker of muscle ageing. PLoS One 8:. https://doi.org/10.1371/journal.pone.0080520

Honma K, Ruscitto A, Frey AM, Stafford GP, Sharma A (2015) Sialic acid transporter NanT participates in Tannerella forsythia biofilm formation and survival on epithelial cells. Microb Pathog 94:12–20. https://doi.org/10.1016/j.micpath.2015.08.012

Kleikamp HBC, Lin YM, McMillan DGG, Geelhoed JS, Naus-Wiezer SNH, van Baarlen P, Saha C, Louwen R, Sorokin DY, van Loosdrecht MCM, Pabst M (2020a) Tackling the chemical diversity of microbial nonulosonic acids – a universal large-scale survey approach. Chem Sci. https://doi.org/10.1039/C9SC06406K

Kleikamp HBC, Pronk M, Tugui C, Guedes da Silva L, Abbas B, Mei Lin Y, van Loosdrecht MCM, Pabst M (2020b) Quantitative profiling of microbial communities by 2 de novo metaproteomics. bioRxiv 2020:2020.08.16.252924. https://doi.org/10.1101/2020.08.16.252924

Knibbs RN, Goldstein IJ, Ratcliffe RM, Shibuya N (1991) Characterization of the carbohydrate binding specificity of the leukoagglutinating lectin from Maackia amurensis. Comparison with other sialic acid-specific lectins. J Biol Chem 266:83–88

Knirel YA, Shashkov AS, Tsvetkov YE, Jansson PE, Zähringer U (2003) 5,7-Diamino-3,5,7,9-tetradeoxynon-2-ulosonic acids in bacterial glycopolymers: Chemistry and biochemistry. Adv Carbohydr Chem Biochem 58:371–417. https://doi.org/10.1016/S0065-2318(03)58007-6

Lambre CR, Kazatchkine MD, Maillet F, Thibon M (1982) Guinea pig erythrocytes, after their contact with influenza virus, acquire the ability to activate the human alternative complement pathway through virus-induced desialation of the cells. J Immunol 128:629–634

Lewis AL, Desa N, Hansen EE, Knirel YA, Gordon JI, Gagneux P, Nizet V, Varki A (2009) Innovations in host and microbial sialic acid biosynthesis revealed by phylogenomic prediction of nonulosonic acid structure. Proc Natl Acad Sci U S A 106:13552–13557. https://doi.org/10.1073/pnas.0902431106

Mainstone CP, Parr W (2002) Phosphorus in rivers - Ecology and management. Sci Total Environ 282–283:25–47. https://doi.org/10.1016/S0048-9697(01)00937-8

Neu T, Kuhlicke U (2017) Fluorescence Lectin Bar-Coding of Glycoconjugates in the Extracellular Matrix of Biofilm and Bioaggregate Forming Microorganisms. Microorganisms 5:5. https://doi.org/10.3390/microorganisms5010005

Neu TR, Lawrence JR (2016) Laser Microscopy for the Study of Biofilms: Issues and Options. In: Romani AM, Guasch H, Balaguer MD (eds) Aquatic Biofilms: Ecology, Water Quality and Wastewater Treatment. Caister Academic Press, Norfolk, pp 29–46

Neu TR, Lawrence JR (2017) The extracellular matrix - an intractable part of biofilm systems. In: Flemming H-C, Neu TR, Wingender J (eds) The perfect slime - Microbial extracellular polymeric substances (EPS). IWA Publishing, London, pp 25–60

Oehmen A, Lopez-Vazquez CM, Carvalho G, Reis MAM, van Loosdrecht MCM (2010) Modelling the population dynamics and metabolic diversity of organisms relevant in anaerobic/anoxic/aerobic enhanced biological phosphorus removal processes. Water Res 44:4473–4486. https://doi.org/10.1016/j.watres.2010.06.017

Oyserman BO, Noguera DR, del Rio TG, Tringe SG, McMahon KD (2016) Metatranscriptomic insights on gene expression and regulatory controls in Candidatus Accumulibacter phosphatis. ISME J 10:810–22. https://doi.org/10.1038/ismej.2015.155

Petit D, Teppa E, Cenci U, Ball S, Harduin-Lepers A (2018) Reconstruction of the sialylation pathway in the ancestor of eukaryotes. Sci REPORTS | 8:2946. https://doi.org/10.1038/s41598-018-20920-1

Ravindranath MH, Higa HH, Cooper EL, Paulson JC (1985) Purification and characterization of an O-acetylsialic acid-specific lectin from a marine crab Cancer antennarius. J Biol Chem 260:8850–6

Rubio-Rincón FJ, Weissbrodt DG, Lopez-Vazquez CM, Welles L, Abbas B, Albertsen M, Nielsen PH, van Loosdrecht MCM, Brdjanovic D (2019) “Candidatus Accumulibacter delftensis”: A clade IC novel polyphosphate-accumulating organism without denitrifying activity on nitrate. Water Res 161:136–151. https://doi.org/10.1016/J.WATRES.2019.03.053

Schnaar RL, Gerardy-Schahn R, Hildebrandt H (2014) Sialic acids in the brain: Gangliosides and polysialic acid in nervous system development, stability, disease, and regeneration. Physiol Rev 94:461–518. https://doi.org/10.1152/physrev.00033.2013

Seviour RJ, Mino T, Onuki M (2003) The microbiology of biological phosphorus removal in activated sludge systems. FEMS Microbiol. Rev. 27:99–127

Seviour T, Derlon N, Dueholm MS, Flemming H-C, Girbal-Neuhauser E, Horn H, Kjelleberg S, van Loosdrecht MCM, Lotti T, Malpei MF, Nerenberg R, Neu TR, Paul E, Yu H, Lin Y (2019) Extracellular polymeric substances of biofilms: Suffering from an identity crisis. Water Res 151:1–7. https://doi.org/10.1016/J.WATRES.2018.11.020

Shibuya N, Goldstein IJ, Broekaert WF, Nsimba-Lubaki M, Peeters B, Peumans WJ (1987) The elderberry (Sambucus nigra L.) bark lectin recognizes the Neu5Ac(alpha 2-6)Gal/GalNAc sequence. J Biol Chem 262:1596–1601

Soares RMA, Rosangela RM, Alviano DS, Angluster J, Alviano CS, Travassos LR (2000) Identification of sialic acids on the cell surface of Candida albicans. Biochim Biophys Acta - Gen Subj 1474:262–268. https://doi.org/10.1016/S0304-4165(00)00003-9

Song X, Yu H, Chen X, Lasanajak Y, Tappert MM, Air GM, Tiwari VK, Cao H, Chokhawala HA, Zheng H, Cummings RD, Smith DF (2011) A sialylated glycan microarray reveals novel interactions of modified sialic acids with proteins and viruses. J Biol Chem 286:31610–31622. https://doi.org/10.1074/jbc.M111.274217

Swofford DL (2001) PAUP*: Phylogenetic Analysis Using Parsimony (and other methods) 4.0.b5

Thompson JD, Gibson TJ, Higgins DG (2003) Multiple Sequence Alignment Using ClustalW and ClustalX. Curr Protoc Bioinforma 00:2.3.1-2.3.22. https://doi.org/10.1002/0471250953.bi0203s00

Traving C, Schauer R (1998) Structure, function and metabolism of sialic acids. Cell. Mol. Life Sci. 54:1330–1349

Varki A, Schnaar RL, Schauer R (2017) Sialic Acids and Other Nonulosonic Acids. In: Varki A (ed) Essentials of Glycobiology, 3rd edn. Cold Spring Harbor Laboratory Press, Cold Spring Harbor (NY), pp 179–195

Watson SJ, Needoba JA, Peterson TD (2019) Widespread detection of *Candidatus* Accumulibacter phosphatis, a polyphosphate-accumulating organism, in sediments of the Columbia River estuary. Environ Microbiol 21:1369–1382. https://doi.org/10.1111/1462-2920.14576

Weissbrodt DG, Neu TR, Kuhlicke U, Rappaz Y, Holliger C (2013) Assessment of bacterial and structural dynamics in aerobic granular biofilms. Front Microbiol 4:. https://doi.org/10.3389/fmicb.2013.00175

Zilles JL, Peccia J, Kim MW, Hung CH, Noguera DR (2002) Involvement of Rhodocyclus-related organisms in phosphorus removal in full-scale wastewater treatment plants. Appl Environ Microbiol 68:2763–2769. https://doi.org/10.1128/AEM.68.6.2763-2769.2002

