## Supplementary figures and images for "Production of nonulosonic acids in the extracellular polymeric substances of “ *Candidatus* Accumulibacter phosphatis”"

### Supplementary Figure 1

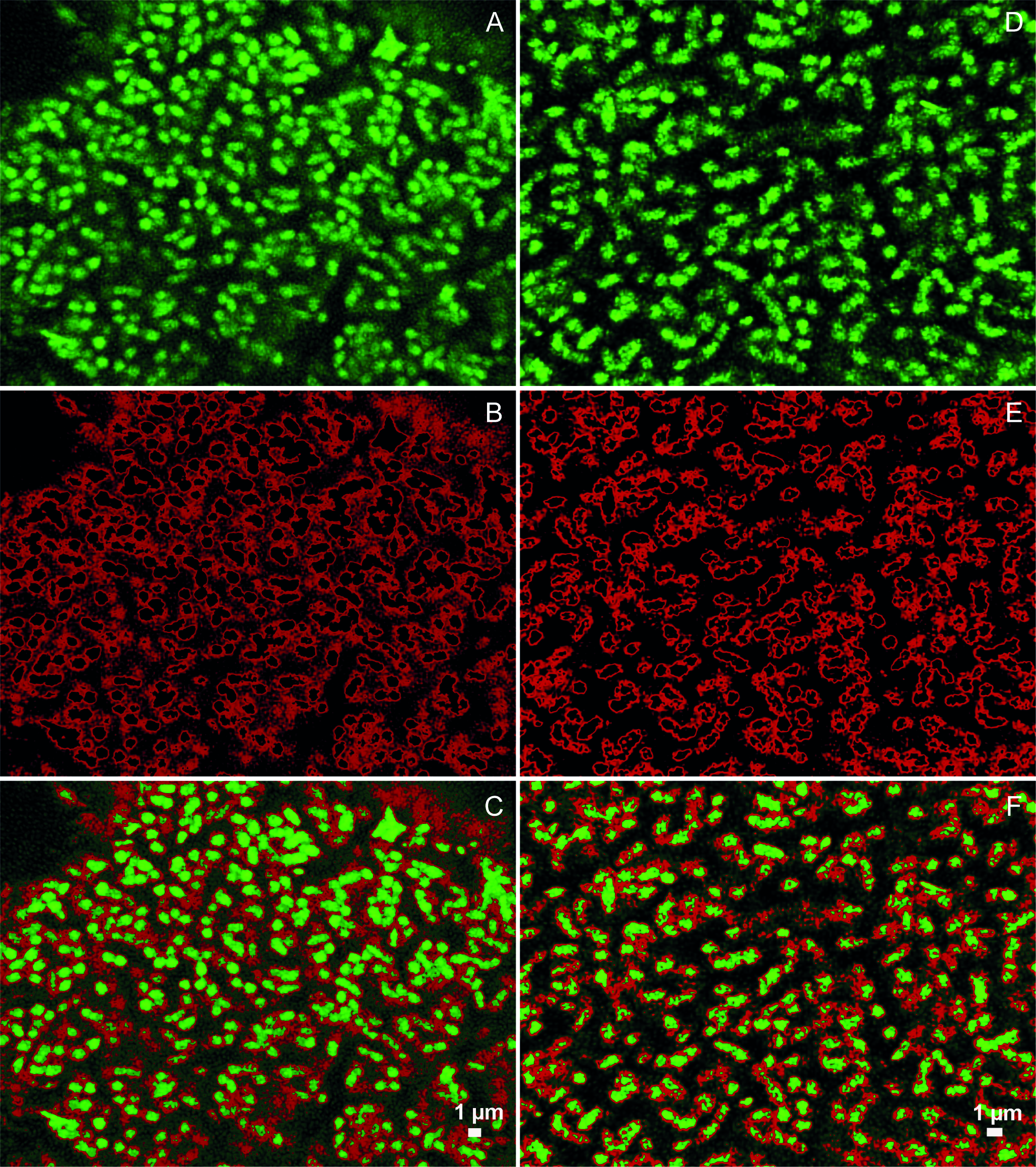
