## Supplementary Materials for "Production of nonulosonic acids in the extracellular polymeric substances of “ *Candidatus* Accumulibacter phosphatis”"

### Supplementary methods

#### Shotgun proteomic analysis

Briefly, biomass material was disrupted using beads beating in B-PER reagent (Thermo Scientific)/TEAB buffer (50 mM TEAB, 1% (w/w) NaDOC, adjusted to pH 8.0) buffer. The cell debris was further pelleted and the proteins were precipitated using ice cold acetone. The protein pellet was redissolved in 200 mM ammonium bicarbonate, reduced using DTT and alkylated using iodoacetamide and digested using sequencing grade Trypsin (Promega). An aliquot of approx. 100 ng protein digest was further analyzed by a one dimensional shotgun proteomics approach (Köcher *et al.*, 2012) using an ESAY nano LC 1200 coupled to a QE plus Orbitrap mass spectrometer (Thermo Fisher Scientific, US). The flow rate was maintained at 300 nL/min over a linear gradient from 5% to 30% solvent B over 85 minutes, and finally to 75% B over 25 minutes. Solvent A was H2O containing 0.1% formic acid, and solvent B consisted of 80% acetonitrile in H2O and 0.1% formic acid. The Orbitrap was operated in data depended acquisition mode acquiring peptide signals form 400-1200 m/z at 70K resolution, where the top 10 signals fragmented using a NCE of 30. Raw data were analyzed using PEAKS Studio 8.5 (Bioinformatics Solutions Inc., Canada) allowing 20 ppm parent ion and 0.02 Da fragment mass error tolerance. Search conditions further included considering 3 missed cleavages, carbamidomethylation as fixed and methionine oxidation and N/Q deamidation as variable modifications. Data were matched against a global “Ca. Accumulibacter phosphatis” database (Uniprot, Date, Tax ID 327159). Peptide search included the GPM crap contaminant database and a decoy fusion for determining false discovery rates. Peptide spectrum matches were filtered against 1% false discovery rate (FDR) and protein identifications with 2 or more unique peptides were considered as significant.

#### Enzymatic quantification

The Sialic Acid Quantitation Kit (Sigma-Aldrich, USA) was used to estimate the content of sialic acids (Neu5Ac as model one) in the enriched “*Ca*. Accumulibacter” biomass. The whole cell assay described in the kit was conducted. Fresh samples were washed with 20 mM Tris-HCl buffer (pH 7.5) and resuspended in 80 μL of demineralized water. Then 20 μL of sialidase buffer and 1 μL of α(2🡪3,6,8,9)-neuraminidase were added. This enzyme releases α-2,3-, α-2,6-, α-2,8-, and α-2,9-linked *N*-acetylneuraminic acid from complex carbohydrates. Samples were incubated overnight at 37°C. After incubation, supernatants were collected, volumes were adjusted to 980 μL with Tris-HCl buffer, 20 μL of 0.01 M β-NADH solution was added and the absorbance at 340 nm was measured. Afterwards, 1 μL of *N*-acetylneuraminic acid aldolase and 1 μL of lactic dehydrogenase were added to each sample, and they were incubated at 37 °C for 1 h. Absorbance at 340 nm was measured again after incubation. Sialic acid concentration was calculated using a calibration line of Neu5Ac provided in the kit.

#### References

Köcher, T., Pichler, P., Swart, R., and Mechtler, K. (2012) Analysis of protein mixtures from whole-cell extracts by single-run nanolc-ms/ms using ultralong gradients. *Nat Protoc* **7**: 882–890.
